# PHC2-Dependent PRC1 Assemblies Stabilize Motor Neuron *Hox* Codes

**DOI:** 10.64898/2026.10.06.757095

**Authors:** Alexander K. Miller, Ayana Sawai, Anne E. Cavanagh, Jeremy S. Dasen

## Abstract

Polycomb Repressive Complexes PRC1 and PRC2 establish and maintain chromatin architectures critical for cell-type specific gene expression. In the developing spinal cord, PRC1 is essential for implementing Hox-dependent transcriptional programs that diversify motor neuron identities. Although PRC1 can assemble into subnuclear condensates, whether these structures contribute to the specification and maintenance of neuronal subtype fates remains unclear. Using chick and mouse models, we show that canonical PRC1 components - BMI1, PHC2, CBX2, RING1B - coalesce into subnuclear assemblies as neural progenitors differentiate into motor neurons.

Disrupting PHC2 polymerization in either neural progenitors or postmitotic neurons dismantles BMI1-containing PRC1 assemblies, triggering derepression of caudal *Hox* genes and a cell autonomous loss of segment-specific motor neuron fates. Mouse mutants lacking the core PRC1 subunit *Ring1* fail to form BMI1-containing assemblies, despite intact PHC2 and CBX2 polymers. In contrast, motor neuron-restricted deletion of core PRC2 subunits (*Ezh1/2*) depletes H3K27me3 histone marks but retains PRC1 assemblies and *Hox* boundaries. Together, these findings indicate that PHC2- dependent PRC1 assemblies stabilize *Hox* transcriptional boundaries and motor neuron segmental identities.

**Summary statement:** Polymerization of the PRC1 subunit PHC2 is essential for the formation of BMI1- containing subnuclear assemblies which establish and maintain *Hox* codes in spinal motor neurons.

## INTRODUCTION

Generation of cell type diversity in the nervous system relies on the coordinated action of spatial and temporal patterning factors coupled to dynamic changes in chromatin structure (Cardona et al., 2025, Sen, 2023). In the developing neural tube, morphogens acting along the dorsoventral (DV) axis specify cardinal classes of spinal neurons (Sagner and Briscoe, 2019). These pathways operate in conjunction with temporal patterning factors to establish distinct transcription factor (TF) codes in both neural progenitors and postmitotic neurons (Zhang et al., 2025). The diversification of neuronal classes in the spinal cord and hindbrain into segmentally restricted subtypes depends on chromosomally arrayed *Hox* genes (Philippidou and Dasen, 2013, Parker and Krumlauf, 2020). In contrast to DV patterning - where distinct sets of TFs define progenitor and postmitotic neuronal identities - *Hox* expression profiles are often maintained throughout both phases of maturation (Bel-Vialar et al., 2002, Dasen et al., 2003, Graham et al., 1991, Liu et al., 2001). How *Hox*-dependent rostrocaudal positional identities are established and stably maintained in developing neurons remains poorly understood.

One class of chromatin regulators governing *Hox* expression is the Polycomb Group (PcG) family (Miller and Dasen, 2024, Gentile and Kmita, 2020). PcG proteins assemble into two evolutionarily conserved Polycomb Repressive Complexes, PRC1 and PRC2, which regulate gene expression through histone modifications and regulation of chromatin structure (Schuettengruber et al., 2007, Simon and Kingston, 2009). In the canonical pathway, PRC2 catalyzes trimethylation of histone H3 at lysine 27 (H3K27me3), serving as a recruitment mark for PRC1 (Margueron and Reinberg, 2011). PRC1 contains subunits capable of compacting chromatin and ubiquitinating histone H2A (Blackledge et al., 2015). During rostrocaudal patterning, *Hox* clusters are initially broadly repressed by PcG activity, and H3K27me3 marks are progressively removed from *Hox* genes as the body axis extends (Boyer et al., 2006, Soshnikova and Duboule, 2009). In spinal progenitors, retinoic acid (RA) from the paraxial mesoderm promotes removal of PRC2 marks and expression of rostral *Hox* genes (*Hox1-Hox5* paralogs), while fibroblast growth factor (FGF) and Wnt gradients derepress more caudal *Hox* genes (*Hox6-Hox10*) (Bel-Vialar et al., 2002, Liu et al., 2001, Mazzoni et al., 2013, Dasen et al., 2003). Removal of PcG-repression enables *Hox* expression which in turn specifies the segmental identities of multiple neuronal classes, including spinal motor neurons (MNs) (Dasen et al., 2005, Sawai et al., 2022, Kratsios et al., 2024).

Maintenance of appropriate *Hox* profiles in spinal MNs depends on PRC1 function (Golden and Dasen, 2012, Sawai et al., 2022). Canonical PRC1 contains four core subunits – a Chromobox (CBX) protein, a Polyhomeotic Homolog (PHC) protein, a Polycomb Group Ring Finger protein (e.g. BMI1), and a RING1 protein. BMI1, PHC, CBX, and RING1 proteins can assemble into subnuclear structures known as PcG bodies (Pirrotta and Li, 2012). The formation of these PRC1 assemblies is essential for maintenance of transcriptional repression (Kishi and Gotoh, 2018). PRC1 condensates formed through polymerization of CBX and PHC proteins promote chromatin compaction, restrict promoter accessibility, and exclude RNA polymerase II from target loci (Niekamp et al., 2024, Plys et al., 2019, Seif et al., 2020, Eskeland et al., 2010). Disruption of CBX- or PHC-mediated polymerization leads to *Hox* gene derepression, homeotic transformations, and widespread transcriptional misregulation (Isono et al., 2013, Isono et al., 2005, Lau et al., 2017, Cheutin and Cavalli, 2018).

Genetic studies in mice indicate that PRC1 and PRC2 play distinct roles in regulating *Hox*-dependent transcriptional programs in spinal MNs. Deletion of the core PRC1 subunit *Ring1* results in marked changes in *Hox* expression and MN subtype specification (Sawai et al., 2022). In *Ring1* mutant mice, the normal *Hox* boundaries in MNs are scrambled, leading to a loss of all Hox-dependent programs. In contrast, removal of PRC2 activity from MN progenitors has little impact on *Hox* expression during embryonic development (Golden and Dasen, 2012, Sawai et al., 2022). These observations suggest that PRC1 plays a dominant role in maintaining *Hox* profiles in postmitotic MNs, but the mechanisms by which PRC1 enforces this repression, particularly when PRC2 activity is diminished, remain unclear. One possibility is that PRC1-dependent subnuclear assemblies stabilize *Hox* repression independently of ongoing PRC2 activity.

Here, we investigate the formation and function of PHC-dependent PRC1 assemblies during spinal MN differentiation. We show that PcG bodies assemble and increase in size as neural progenitors differentiate into MNs. Disruption of PHC2 polymerization in chick embryos attenuates PRC1 assembly formation, leading to derepression of caudal *Hox* genes, erosion of MN *Hox* codes, and loss of Hox-dependent MN subtypes.

Genetic removal of *Ring1* from MNs in mice abolishes BMI1-containing assemblies while leaving CBX and PHC2 assemblies intact. By contrast, disruption of PRC2 activity has no detectable effect on PcG body maintenance or *Hox* repression. Together, these findings demonstrate that canonical PRC1 maintains Hox-dependent spinal MN positional identities through PHC-dependent subnuclear assemblies.

## RESULTS

### Subnuclear localization of PRC1 components during motor neuron differentiation

To assess a potential role for PRC1 assemblies in establishing and maintaining *Hox* profiles in MNs, we first performed an analysis of the distribution of PRC1 proteins within the developing spinal cord. We examined the distribution of 4 canonical PRC1 components (BMI1, PHC2, CBX2, and RING1B) in Hamburger & Hamilton (HH) stage 27 chick embryos. We selected these four PRC1 proteins based on bulk RNAseq data from MNs isolated from E12.5 mouse spinal cord indicating prominent expression of these genes relative to other PRC1 subunits and paralogs (**Fig. S1A**) (Sawai et al., 2022). Histological analyses showed that BMI1, PHC2, CBX2, and RING1B proteins were present within both neural progenitors (NPs) and postmitotic HB9^+^ MNs (**Fig. S1B**). Expression of BMI1, PHC2, and CBX2 was similar in NPs and MNs, while RING1B was enriched in MNs (**Fig. 1A-D**, **S1B**). Similar patterns of BMI1, PHC2, CBX2, and RING1B protein expression were observed in E12.5 mouse spinal cord (**Fig. S1C**).

**Fig. 1:**
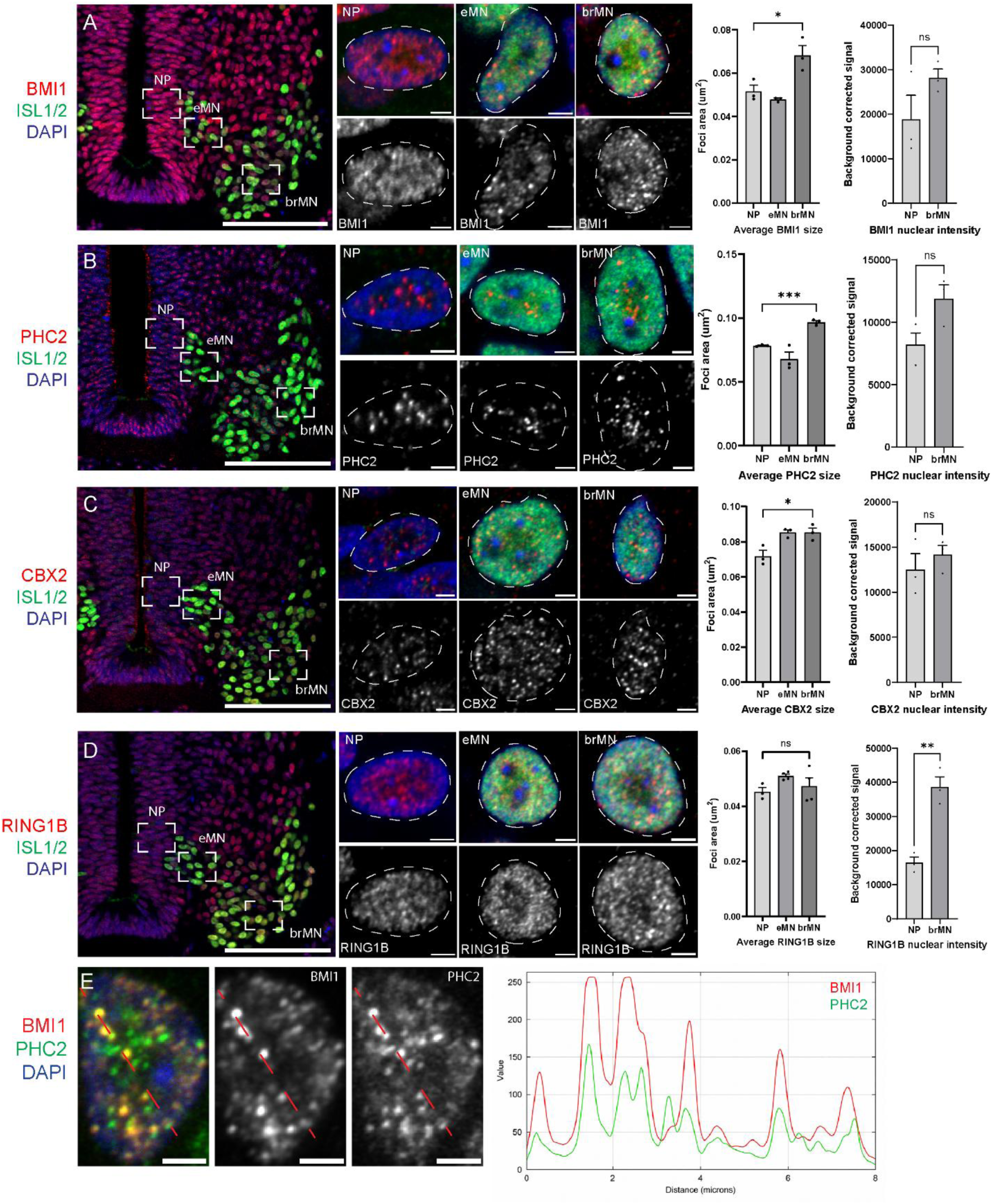
PRC1 subunit assemblies during motor neuron differentiation. (A-D) High resolution immunofluorescent images of core PRC1 components within neural progenitor (NP), early motor neuron (eMN), and brachial motor neuron (brMN) nuclei. White boxes mark regions used for high-resolution imaging. Right panels show comparison of puncta sizes in NP, eMN, and brMN populations and nuclear signal intensities in NP and brMN. Data from n=3 independent embryos. Bar graphs show mean ± SEM for n=3-4 embryos, 3 sections/embryo. *p < 0.05, **p < 0.01, ***p < 0.001, unpaired t-test. ns=not significant. Scale bars for spinal cord images, 100µm. Scale bars for nuclei, 2µm. (E) Co-localization of BMI1 and PHC2 in MNs. Plot on right shows BMI1 and PHC2 intensities along a scan line in images on left. Scale bars, 2µm.

As BMI1, PHC, CBX, and RING1 are major constituents of PcG bodies, we used high resolution (Airyscan) confocal microscopy to assess PRC1 subnuclear assemblies during the progression of NPs to MNs. We compared the distribution of PRC1 proteins in NPs, early postmitotic motor neurons (eMNs), and post-migratory motor neurons in brachial segments (brMNs). Within brMNs, BMI1 localized to discrete subnuclear foci, whereas at earlier stages its distribution was more diffuse (**Fig. 1A**). We observed a similar pattern of focalization for PHC2 and CBX2 (**Fig. 1B-C**), whereas RING1B staining was more diffuse throughout differentiation (**Fig. 1D**). Comparison of PRC1 subunit localization between progenitors and brMNs showed an increase in average puncta size after differentiation (**Fig. 1B-E**, **S1D**). Co-staining for BMI1 and PHC2 further revealed co-localization within subnuclear puncta of MNs (Pearson’s correlation coefficient r=0.71), consistent with the presence of PRC1 assemblies (**Fig. 1E**). These observations indicate that PRC1 proteins become increasingly concentrated in larger subnuclear assemblies as neural progenitors mature into postmitotic MNs.

### Inhibition of PHC2 polymerization disrupts PcG body formation in motor neurons

Our analyses of canonical PRC1 show that specific subunits are present as subnuclear assemblies in spinal MNs. The significance of PRC1 assembly formation for the maintenance of transcriptional repression in spinal MNs is unclear. Given that the pattern of *Hox* expression in spinal MNs is resistant to genetic depletion of PRC2 function (Sawai et al., 2022), we sought to address which components of PRC1 might be necessary for maintenance of PcG bodies and *Hox* repression. Previous studies indicate that the PRC1 component PHC polymerizes and is essential for the formation of condensed chromatin regions in both flies and mammals (Isono et al., 2005, Seif et al., 2020). We therefore considered the possibility that PHC-dependent polymerization is involved in the maintenance of *Hox* profiles in spinal MNs.

To test the role of PHC polymerization in PRC1-dependent repression, we expressed a dominant negative PHC2 (*dnPhc2*) construct, which contains a point mutation (L307R) that inhibits polymerization through blocking association between PHC2 SAM-domains (**Fig. 2A**) (Isono et al., 2005). Because of potential functional compensation between the three *Phc* genes, we chose a dominant negative strategy to assess the role of these proteins. The *dnPhc2* construct was expressed in the neural tube of stage 15 chick embryos under a constitutive promoter (*pCAGGs*) containing an ires-driven nuclear *GFP* (**Fig. 2A**). We also compared the effects of *dnPhc2* expression with unmodified *Phc2*. Embryos were allowed to develop for 2-3 days (HH25-31) until the time of spinal MN differentiation. To quantify the level of exogenous *Phc2* expression, fluorescent intensities of PHC2+ nuclei were measured following electroporation of *dnPhc2*. Within GFP^+^ nuclei, dnPHC2 expression was increased ∼5-fold relative to endogenous protein levels, validating our ability to express *dnPhc2* within the developing spinal cord without impacting cell viability (**Fig. 2B**). Importantly, two markers of core MN identity, HB9 and ISL1/2 (Thaler et al., 2004, Arber et al., 1999), were not affected after expression of *dnPhc2* or unmodified *Phc2* (**Fig. 2G, H**).

**Fig. 2:**
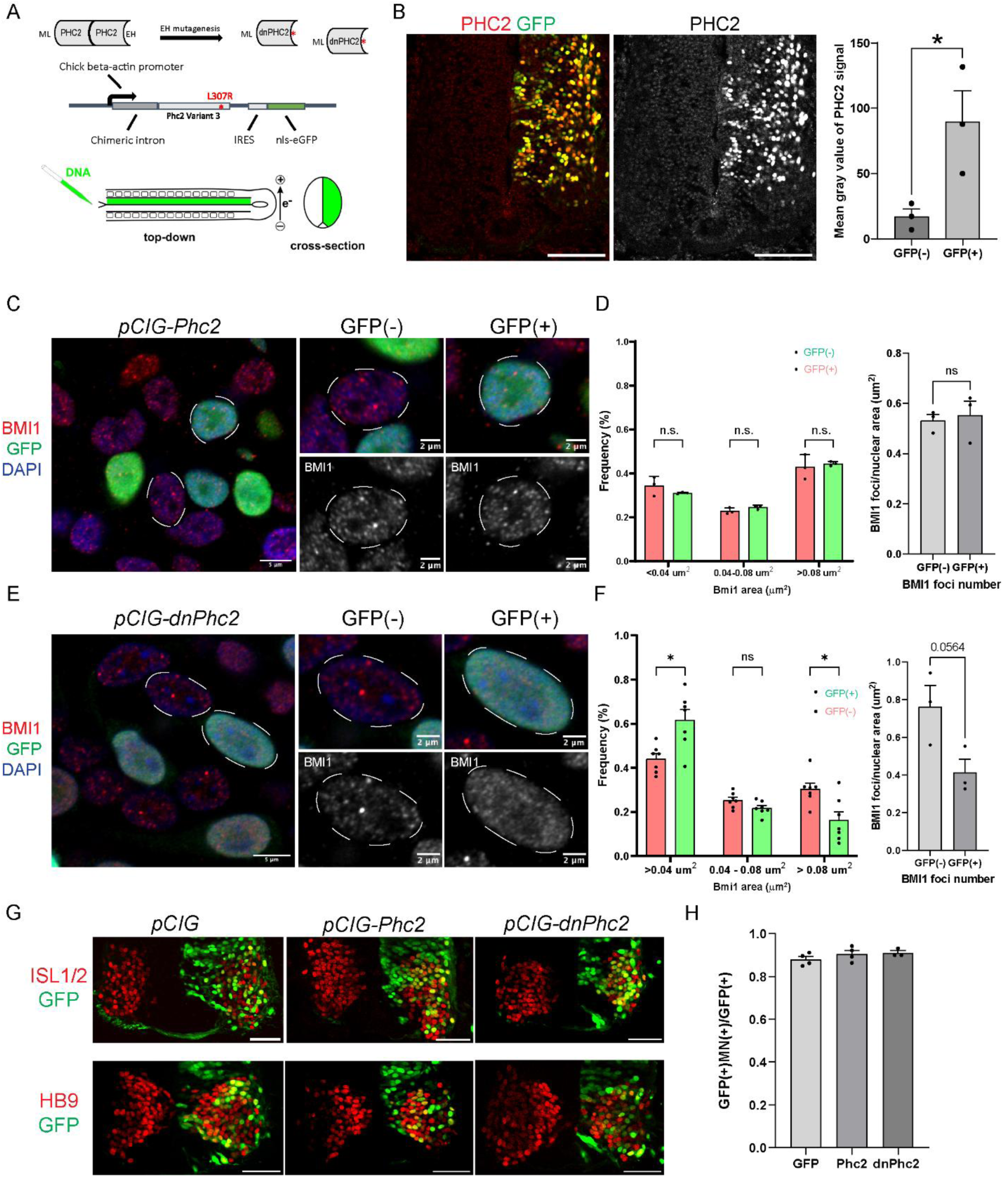
Expression of *dnPhc2* within the neural tube reduces PRC1 assemblies. (A) Diagram of the *dnPhc2* construct and its predicted effect of PHC2-dependent polymerization. Chick electroporation schematic shown below. (B) PHC2 fluorescent intensities in GFP(-) and GFP(+) nuclei. Right side of the spinal cord is electroporated. Bar graphs show mean ± SEM for n=3 embryos, 4 sections/embryo. *p < 0.05, unpaired t-test. Scale bars, 100µm. (C-F) High-resolution Airyscan imaging of BMI1 staining in embryos electroporated with *pCIG-Phc2* or *pCIG-dnPhc2* constructs. Right panels show the distribution of BMI1 puncta sizes binned as small (<0.04 µm^2^), medium (0.04-0.08 µm^2^), and large (>0.08 µm^2^) and number of foci per area. Bar graphs show mean ± SEM for n=3-5 embryos, 3 sections/embryo. *p < 0.05, unpaired t-test. Scale bar sizes shown in panels. (G) Expression of the general MN markers ISl1/2 and HB9 after electroporation of *pCIG*, *pCIG-Phc2* and *pCIG-dnPhc2* constructs. Scale bars, 50 µm. (H) Quantification of the fraction of electroporated GFP+ cells expressing ISL1/2 and HB9 after electroporation of each construct. Bar graphs show mean ± SEM for n=3-4 embryos/condition, 3 sections/embryo. ns=not significant, unpaired t-test.

To quantify possible effects on PcG bodies following *dnPhc2* expression, we performed high-resolution imaging of the subnuclear distribution of the PRC1 protein BMI1 in brachial segments (**Fig. 2C-F**). We chose to focus on BMI1 for these analyses for several reasons. Antibodies against PHC2 also recognize exongenously expressed *dnPhc2*, making PHC2 localization within GFP^+^ nuclei difficult to quantify (**Fig. S2B**).

Although BMI1 possesses no polymerization capacity by itself, its subnuclear clustering is likely a consequence of recruitment to PRC1 assemblies, via direct interactions with PHC proteins (Gray et al., 2016). Thus, a change in the subnuclear distribution of BMI1 would reflect a broader impact on PRC1 assembly formation.

We quantified a potential reduction in BMI1-containing subnuclear assemblies by assessing the distribution of puncta densities and sizes in GFP^+^ and GFP^-^ nuclei in *dnPhc2* and control *Phc2* electroporated embryos (**Fig. 2C-F**; **S2A, B**). To simplify these analyses, we grouped nuclear puncta in three area bins (< 0.04 µm^2^, 0.04 µm^2^ - 0.08 µm^2^ , and > 0.08 µm^2^). We observed a reduction in the number of large BMI1 foci (> 0.08 µm^2^ in area) in GFP^+^ nuclei, and an increase in smaller and more diffuse foci (< 0.04 µm^2^ in area) (**Fig. 2E,F**), in *dnPhc2* embryos consistent with disruption of PHC2- dependent PRC1 assembly. By contrast, expression of unmodified *Phc2* had no impact on the size or distribution of BMI1 or PHC2 containing nuclear puncta (**Fig. 2C,D**, **S2C**). The size of CBX2-containing puncta was also unaffected in *dnPhc2* expressing neurons (**Fig. S2D**). The persistence of CBX puncta following disruption of PHC2 polymerization is consistent with evidence that CBX proteins can initiate subnuclear clustering, while PHC2 polymerization plays a critical role in stabilizing and organizing higher-order PRC1 condensates (Niekamp et al., 2024).These results show that expression of *dnPhc2* can reduce the size of BMI1-containing PRC1 assemblies *in vivo*, without affecting MN class specification, allowing us to examine the requirement of PHC2- dependent polymerization in sustaining *Hox* boundaries in MNs.

### Blocking PHC2 polymerization results in caudal *Hox* gene derepression

To assess the consequences of disruption on PHC2 polymerization, we examined *Hox* expression within brachial and thoracic segments of the spinal cord. Following expression of *dnPhc2*, we observed a derepression of caudal *Hox* paralogs in rostral segments, exemplified by a pronounced rostral misexpression of HOXC9 in brachial segments (**Fig. 3A**, **S3C**). Comparing HOXC9 immunofluorescent intensities with the average size of BMI1 foci in GFP^+^ nuclei revealed elevated HOXC9 expression in nuclei with a more pronounced reduction in BMI1 foci size (**Fig. 3B-D**). By contrast, HOXC9 expression was not significantly changed following neural tube expression of either unmodified *Phc2* or *GFP* alone (**Fig. 3A**, **S3A**).

**Fig. 3:**
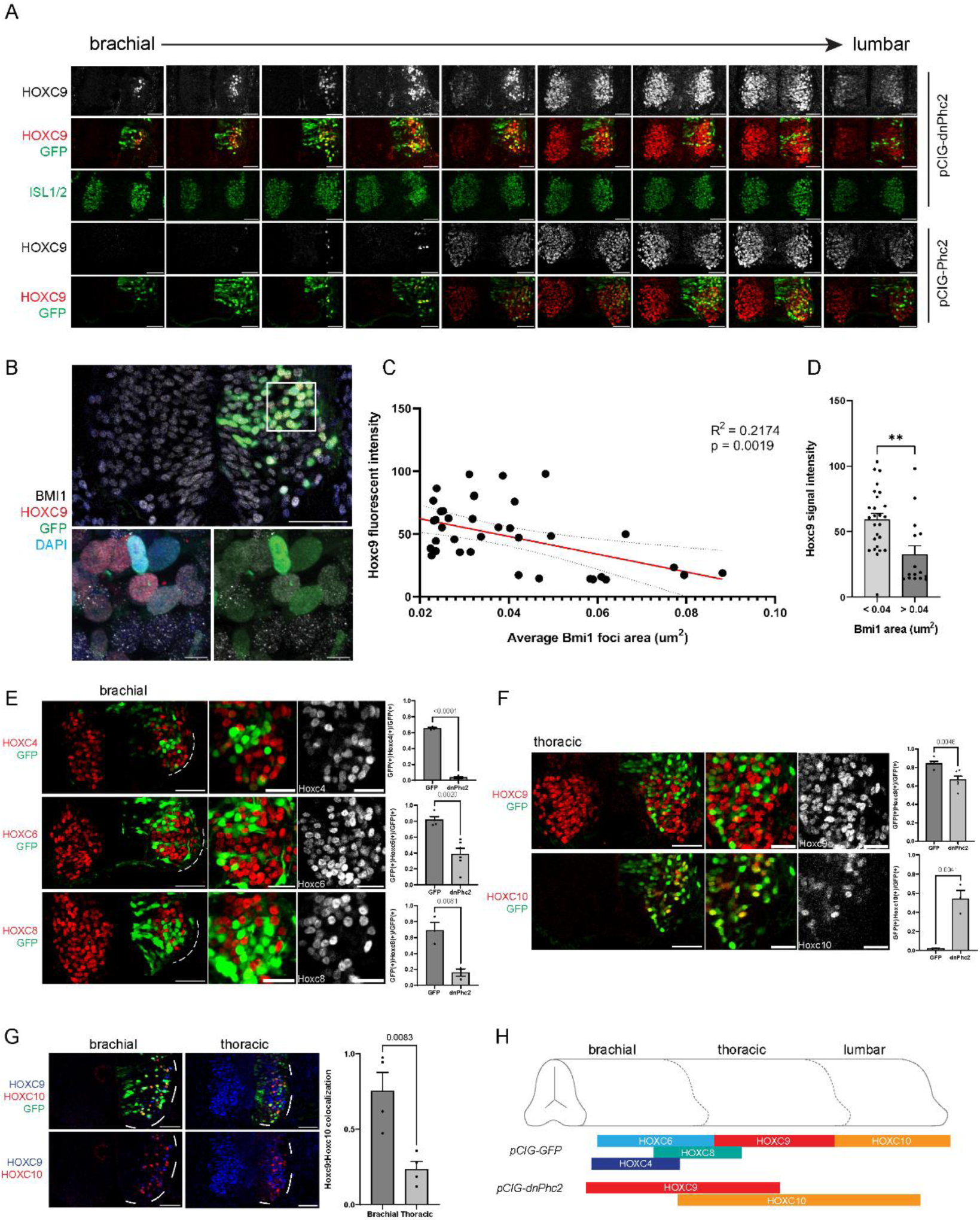
Inhibition of PHC2 polymerization leads to caudal misexpression of *Hox* genes. (A) Serial sections along rostrocaudal axis showing ectopic HOXC9 expression extends into brachial segments after *pCIG-dnPhc2* electroporation. A few ectopic HOXC9 cells are observed near the brachial/thoracic boundary after *pCIG-Phc2* expression. Expression of general MN marker ISL1/2 is unaffected. Right side of the spinal cord is electroporated. Distance between adjacent sections in figure is ∼120 µm. Scale bars, 50 µm (B) Representative images of HOXC9 misexpression within *dnPhc2*-electroporated embryos. Scale bars, 50 µm (low resolution images), 2 µm (high-resolution images). (C) Linear correlation of HOXC9 fluorescent intensity and average BMI polymer sizes. (D) Comparison of ectopic HOXC9 fluorescent intensities between brachial MNs with small (<0.04 µm^2^) versus large (> 0.04 µm^2^) BMI1 puncta. Bar graphs show mean ± SEM. ****p < 0.0001, unpaired t-test. (E) Immunofluorescence of HOX proteins in brachial spinal cord. Quantification shown on right. Bar graphs show mean ± SEM for 3-6 embryos, 12 sections/embryo. **p < 0.01, ****p < 0.0001, unpaired t-test. Scale bars, 50 µm. (F) Immunofluorescence of HOXC9 and HOXC10 in thoracic spinal cord. Quantification of *Hoxc9/10:GFP* colocalization is shown on right. Bar graphs show mean ± SEM for n=3-6 embryos, 12 sections/embryo. **p < 0.01, ****p < 0.0001, unpaired t-test. Scale bars, 50 µm. (G) Expression of HOXC9 and HOXC10 in brachial and thoracic segments. Bar graphs show mean ± SEM for n=4 embryos, 3 sections/embryo. ****p < 0.0001, unpaired t-test. Scale bars, 50 µm. (H) Schematic summarizing HOX misexpression following expression of *dnPhc2*.

The *Hoxc9* gene is normally required for the exclusion of *Hox4-Hox8* paralog expression from thoracic segments, and HOXC9 can repress these *Hox* genes when expressed in brachial segments (Jung et al., 2010, Jung et al., 2014). In agreement with these findings, misexpression of HOXC9 after *dnPhc2* expression led to repression of HOXC4, HOXC6, and HOXC8 in GFP+ nuclei (**Fig. 3E**). Moreover, after *dnPhc2* expression the lumbar-expressed *Hox* gene HOXC10 was detected in brachial segments (**Fig. 3G**, **S3B,C**). As a result, in brachial segments HOXC9 and HOXC10 are co-expressed in spinal MNs (**Fig. 3G**). This result is reminiscent of the Hox co-expression phenotype in mice lacking the core PRC1 component *Ring1* (Sawai et al., 2022).

In thoracic segments expression of *dnPhc2* led to a more pronounced derepression of HOXC10 and reduced expression of HOXC9. Motor neurons that expressed HOXC10 in thoracic segments lacked HOXC9 (**Fig. 3F**). This results likely reflects segment-specific differences in the degree of caudal *Hox* derepression; with the more pronounced derepression of HOXC10 in thoracic segments leading to repression of *Hoxc9*. These findings are reminiscent of the *Hox* expression changes in *Ring1* mutants, which are characterized by both a caudal shift and scrambling of the normal MN Hox codes.

Together these results demonstrate that expression of *dnPhc2* leads to derepression of caudal *Hox* genes in more rostral segments (**Fig. 3H**). This results in the dissolution of Hox boundaries, exemplified by the misexpression of HOXC9 and HOXC10 in brachial and thoracic segments and a repression of more rostral *Hox* genes through cross-repressive interactions.

### Expression of *dnPhc2* results in a loss of *Hox*-dependent motor columns

Disruption of PHC2 polymerization impairs the ability of PRC1 to maintain *Hox* boundaries, resulting in both a caudal shift and intermixing of *Hox* pattern. We next asked how these changes in *Hox* expression affect the programming of MN subtype identities. Spinal MNs segregate into longitudinally-arrayed columns with distinct molecular identities, anatomical positions, and peripheral targets (Philippidou and Dasen, 2013). At brachial levels, specification of the forelimb-innervating lateral motor column (LMC) requires *Hox4-Hox8* genes, while the thoracic-restricted preganglionic motor column (PGC) is dependent on *Hoxc9*. In addition, two axial motor columns, the medial (MMC) and hypaxial motor columns (HMC) appear to be specified independent of *Hox* function (Dasen, 2022).

To assess the consequences of *Hox* misexpression on MN columnar identity, we examined expression of FOXP1, a downstream effector of HOX proteins in both LMC and PGC neurons (Dasen et al., 2008, Rousso et al., 2008). Following expression of *dnPhc2*, FOXP1 expression was markedly reduced in GFP^+^ MNs of brachial and thoracic segments (**Fig. 4A**). A nearly complete loss of FOXP1 in GFP+ MNs suggests both LMC and PGC populations are not specified in *dnPhc2*-expressing MNs.

**Fig. 4:**
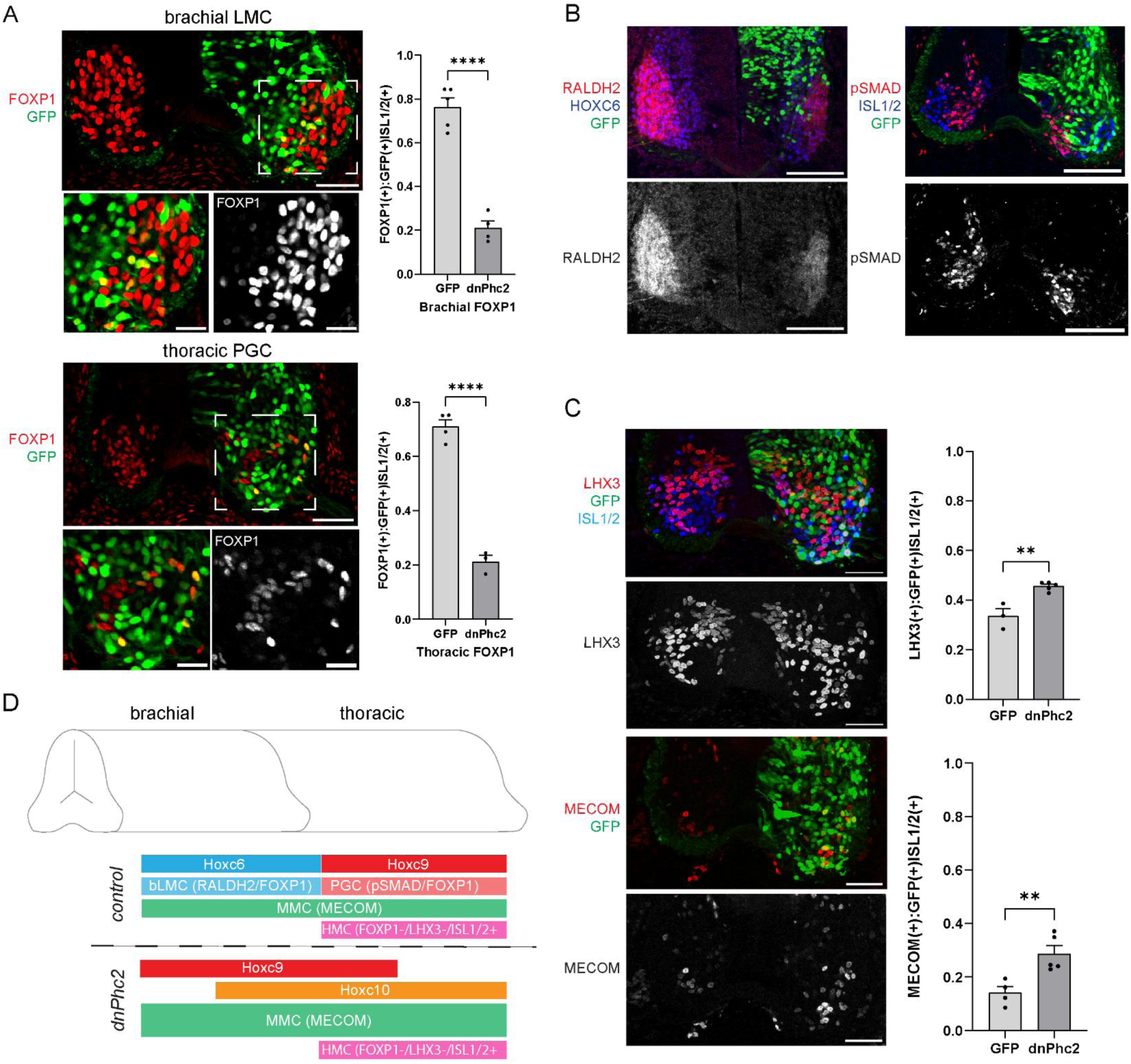
Loss of *Hox-*dependent motor columns after expression of *dnPhc2*. (A) Expression of the LMC and PGC marker FOXP1 in brachial and thoracic spinal cord sections after expression of *dnPhc2*. Right side of the spinal cord is electroporated. Graphs on right show FOXP1 expression between GFP(-) vs GFP(+) cells. Bar graphs show mean ± SEM for n=4 embryos, 3 sections/embryo. ****p < 0.0001. Scale bars, 50 µm (top); 25 µm (bottom). (B) Expression of the LMC-restricted marker RALDH2 and the PGC marker pSMAD are reduced in *dnPhc2* electroporated MNs. (C) Immunofluorescence of markers for *Hox-*independent MNs subtypes, HMC (LHX3/ISL1/2^+^) and MMC (MECOM^+^). Quantification of LHX3^+^ and MECOM^+^ cells between *dnPhc2* electroporated and unelectroporated embryos shown on the right. Bar graphs show mean ± SEM for n=3-5 embryos, 3 sections/embryo. **p < 0.01, unpaired t-test. Scale bars, 50 µm. (D) Schematic summarizing the effects of *dnPhc2* expression on MN organization.

Consistent with this, the brachial LMC marker RALDH2 and the thoracic PGC marker pSMAD were depleted from *dnPhc2*-expressing MNs (**Fig. 4B**), indicating a combined loss of both Hox-dependent MN columnar identities.

We next explored the fates of the remaining MNs after *dnPhc2* expression. We assessed two Hox-independent axial muscle innervating MN subtypes, the medial motor column (MMC) and hypaxial motor column (HMC). For MMC neurons we assessed expression of MECOM^+^ and LHX3^+^ in MNs, while HMC neuron were defined as ISL1^+^ MNs lacking MECOM and FOXP1 expression (Hanley et al., 2016, Tsuchida et al., 1994). After *dnPhc2* expression the number of MECOM^+^ and LHX3^+^ significantly increased in brachial and thoracic spinal cord, while the number of HMC neurons was unchanged (**Fig. 4C**). Overall, the reduction in PHC polymerization, and misexpression of *Hox* transcription factors, result in presumptive LMC and PGC MNs acquiring a *Hox*- independent MMC axial MN fate (**Fig. 4D**).

### Motor neuron-restricted expression of *dnPhc2* results in *Hox* derepression

The increase in the size of PRC1 subnuclear foci during differentiation suggests a continuous reliance on PRC1-mediated repression in postmitotic neurons. The elevated expression of PHC2 in MNs suggests PHC-dependent polymerization plays a critical role. We therefore examined the effects of disrupting PHC2 polymerization on the maintenance of *Hox* boundaries in MNs.

We expressed a *dnPhc2* under control of the *Hb9*-promoter which targets postmitotic spinal MNs (Wichterle et al., 2002). We generated and tested two *dnPhc2* constructs, one consisting of unmutated *Phc2* fused directly to a C-terminal GFP tag, and a second consisting of a direct fusion with the SAM-domain point mutation (**Fig. 5A**). Addition of a GFP tag to the C-terminal end of PHC2 has previously been shown to sterically hinder dimerization and inhibit PHC2-dependent polymerization (Isono et al., 2013).

**Fig. 5:**
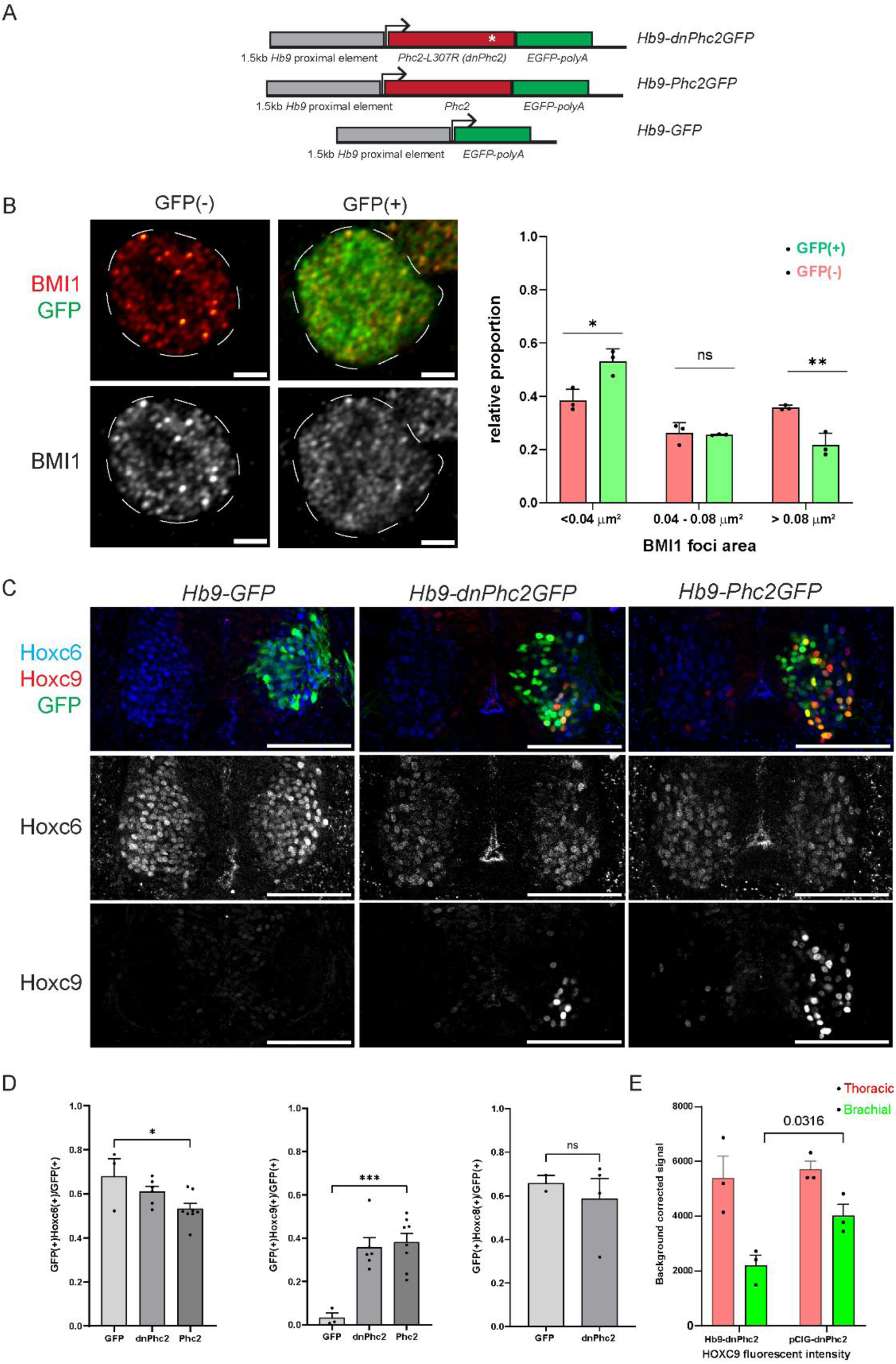
PHC polymerization in postmitotic MNs sustains *Hox* gene repression. (A) Schematics of *Hb9-dnPhc2GFP*, *Hb9-Phc2GFP*, and *Hb9-GFP* constructs. (B) Representative images of BMI1 assemblies in chick MNs after expression of *Hb9-dnPhc2GFP*. Scale bars, 2 µm. Right: the proportion of BMI1 foci sizes binned as small (<0.04 µm^2^), medium (0.04-0.08 µm^2^), or large (>0.08 µm^2^). Graphs show mean ± SEM for n=3 embryos, 3 sections/embryo. *p < 0.05, **p < 0.01, unpaired t-test. (C) Expression of HOXC6 and HOXC9 in electroporated MNs. Right side of spinal cord is electroporated. Scale bars, 100 µm. (D) Quantification of HOXC6 and HOXC9 in brMNs of *Hb9-GFP* (GFP), *Hb9*-*dnPhc2GFP* (dnPhc2), and *Hb9*-*Phc2GFP* (Phc2) electroporated MNs. Bar graphs show mean ± SEM for n=3-6 embryos, 3 sections/embryo. *p < 0.01, **p < 0.01, ***p < 0.001, unpaired t-test. (E) Expression levels of HOXC9 achieved using *pCIG-dnPhc2* and *Hb9-dnPhc2* at brachial levels, compared to HOXC9 expression in control thoracic sections. Bar graphs show mean ± SEM for n=3 embryos, 3 sections/embryo. Unpaired t-test P value shown in figure.

Electroporation of *Hb9*-driven constructs localized *dnPhc2* expression to mature MNs (**Fig. 5C**), allowing us to examine necessity of PHC polymers for the maintenance of *Hox* repression in MNs.

Following expression of *Hb9-dnPhc2GFP* or *Hb9-Phc2GFP*, we observed a significant reduction in the size of BMI1 foci within GFP^+^ nuclei, indicating successful reduction in PRC1 assembly formation (**Fig. 5B**). To determine if PRC1 assemblies are important for maintenance of repression within MNs, we examined *Hox* expression within the brachial segments. Following expression of *Hb9-dnPhc2GFP* or *Hb9-Phc2GFP*, we observed a cell-autonomous derepression of HOXC9, which in turn inhibits the expression of brachial HOXC6 (**Fig. 5C, D**). In contrast to *pCIG-dnPhc2* electroporations, *Hb9*-driven expression did not result in a significant reduction of brachial HOXC8 expression. This difference appears to reflects a reduced level of ectopic HOXC9 expression achieved in the *Hb9* context relative to *pCIG-dnPhc2* conditions (**Fig. 5E**). Nevertheless, these results indicate that PHC2-dependent polymerization remains essential for the repression of *Hox* genes in differentiated MNs.

### PRC1 assembly stability is resistant to attenuation of PRC2 activity

Our findings suggest that PHC2 plays a critical role in the maintenance of spinal *Hox* boundaries. Previously, we observed that combined genetic deletion of two core PRC2 components, *Ehz1* and *Ezh2*, or deletion of *Eed*, from MN progenitors had no discernable impact on *Hox* patterning in embryonic spinal MNs (Sawai et al., 2022). The ability of *dnPhc2* to disrupt *Hox* profiles raises the possibility that PHC-dependent polymers can maintain MN positional identities in the absence of continuous PRC2 function. To further explore this idea, we assessed the formation of PRC1-containing PcG bodies in mice lacking PRC2, through genetic depletion of both *Ezh1* and *Ezh2* from spinal MN progenitors (*Ezh MNΔ* mice). In this strategy, expression of *Olig2::Cre* deletes *Ezh* gene function from MN progenitors, and H3K27me3 depletion is maintained in postmitotic MNs.

Analyses of *Ezh MNΔ* mice at E12.5 revealed that while H3K27me3 was depleted from MNs (**Fig. 6A, C**), the overall patterns of BMI1, PHC2, CBX2, and RING1B protein expression were unchanged (**Fig. 6A**). High resolution imaging of spinal MNs revealed the presence of BMI1, PHC2, CBX2, and RING1B containing nuclear puncta (**Fig. 6B**). The average size of BMI1, CBX2 and PHC2 puncta were unaffected in *Ezh MNΔ* mice (**Fig. 6C**, **S4A**). In mouse RING1B staining was too diffuse to measure accurately.

**Fig. 6:**
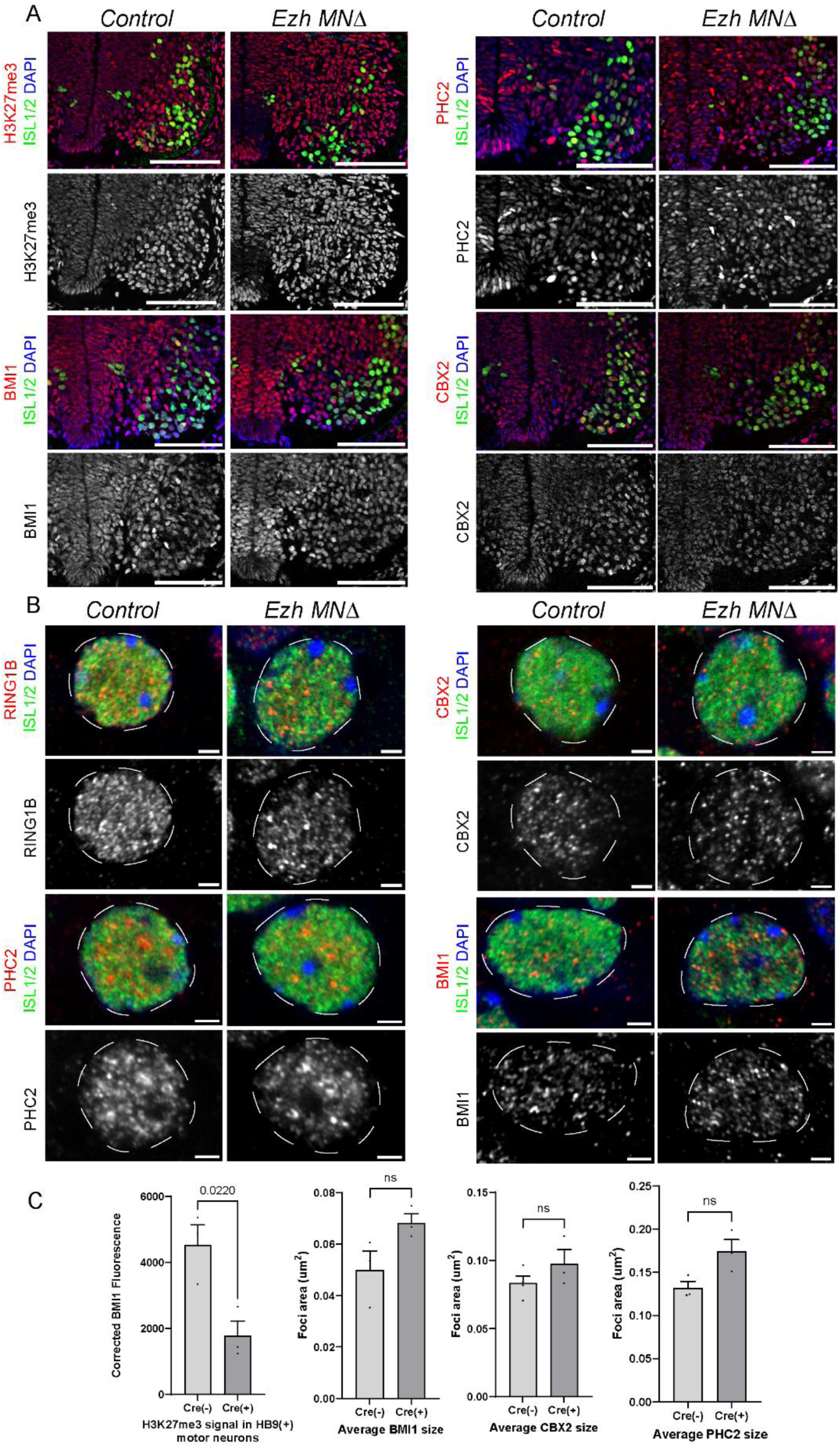
Motor neuron-restricted deletion of *Ezh1/2* does not impair PRC1 assembly. (A) Staining of H3K27me3, PHC2, BMI1, and CBX2 in control (Cre-) or *Ezh1/2 MNΔ* (Cre+) E12.5 mouse embryos. ISL1/2^+^ MNs are shown in green. Scale bars, 100 µm. (B) High-resolution Airyscan images of PRC1 components in control and *Ezh1/2 MNΔ* MNs. Scale bars, 2 µm (C) Quantification of H3K27me3 as well as BMI1, CBX2, and PHC2 puncta in control (Cre-) and *Ezh1/2* mutant (Cre+) MNs. Bar graphs show mean ± SEM for n=3 embryos, 3 sections/embryo.

These results indicate that formation of PRC1 containing PcG bodies is resistant to loss of PRC2 activity.

To further validate these findings, we used chick neural tube electroporation to acutely block PRC2 activity in progenitors. To inhibit the methyltransferase activity of PRC2, we expressed a dominant negative *Ezh2* construct (*dnEzh2*), which lacks the catalytic SET domain and has been shown to reduce H3K27me3 levels and disrupt DV patterning in the neural tube (Akizu et al., 2010). We expressed *dnEzh2-ires-nlsGFP* at HH15 and analyzed embryos at HH25 (**Fig. S5A**). Staining for H3K27me3 revealed a dose-dependent reduction of H3K27me3 in electroporated neurons, with an ∼80% reduction in H3K27me3 signal at the highest plasmid concentration we tested (**Fig. S5C**).

Reduced H3K27me3 had no detectable impact on the domain of *Hoxc9* expression (**Fig. S5B**). Moreover, high-resolution imaging of MNs showed no difference in PRC1 assembly size between GFP^−^ and GFP^+^ cells, as assessed by analyses of BMI1 and PHC2 (**Fig. S5D**). Collectively, these results indicate that PcG body stability is resistant to loss of PRC2 activity and attenuated H3K27me3 levels.

### *Ring1* is required for formation of BMI1-containing PcG bodies in motor neurons

We next investigated the formation of PcG bodies in mice lacking the core PRC1 component *Ring1*. In contrast to PRC2-depleted mouse embryos, genetic removal of *Ring1* has a dramatic impact on *Hox* boundaries and MN subtype specification. While generic features of MNs are specified normally in *Ring1* mutants, they are characterized by increased accessibility and derepression of dozens of TFs, including caudal *Hox* genes, and a complete loss of Hox-dependent MN subtypes (Sawai et al., 2022).

As previous studies have indicated that both PHC and CBX proteins can assemble into polymers *in vitro*, even in the absence of other PRC1 components (Gemeinhardt et al., 2025, Plys et al., 2019), we first examined the presence of PHC2 and CBX2 nuclear puncta in *Ring MNΔ* mice at E12.5. The overall expression patterns of PHC2 and CBX proteins and mRNAs were not affected by deletion of *Ring1* genes (**Fig. 7A, D**).

**Fig. 7.**
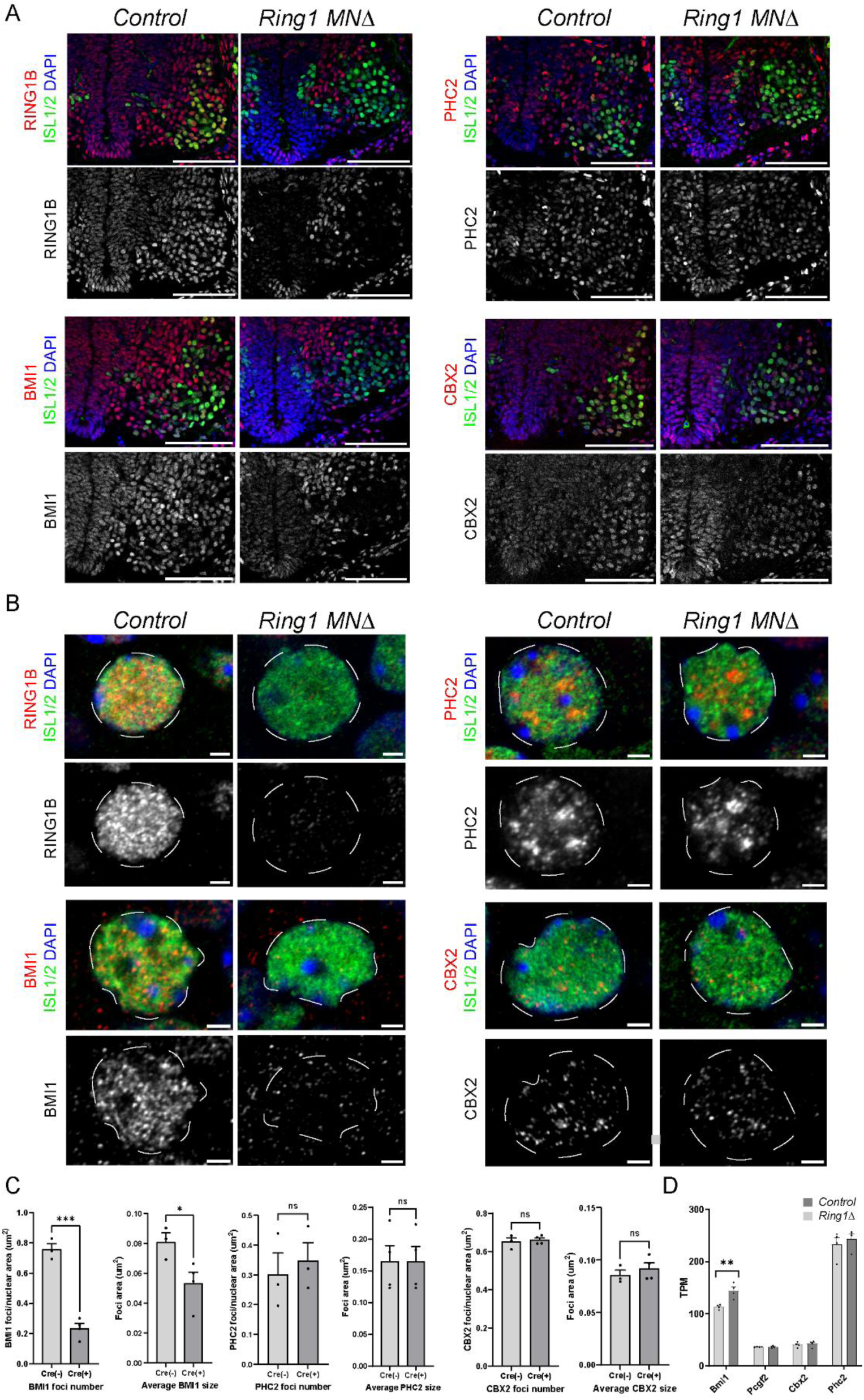
Motor neuron-restricted deletion of *Ring1* genes reduces BMI1-containing PRC1 assemblies. (A) Expression of the RING1B, PHC2, BMI1, and CBX2 in control (Cre-) and *Ring1 MNΔ* (Cre+) e12.5 mouse embryos. ISL1/2^+^ MNs are shown in green. Scale bars, 100 µm. (B) High-resolution Airyscan images of PRC1 components in control and *Ring1* mutant MNs. Scale bars, 2 µm. (C) Quantification of BMI, PHC2, and CBX2 puncta areas and foci number/area in control (Cre-) and *Ring1* mutant (Cre+) MNs. Bar graphs show mean ± SEM for n=3 embryos, 3 sections/embryo. *p < 0.01, ***p < 0.001, unpaired t-test. (D) Bulk RNAseq counts (transcripts per million [TPM]) of *Bmi1*, *Pcgf2*, *Cbx2*, and *Phc2* in control and *Ring1MNΔ* motor neurons. Bar graphs show mean ± SEM for n=4 control and mutant embryos, 3 sections/embryo, **p < 0.01, unpaired t-test.

Moreover, the number and size of PHC2 and CBX2 puncta were unaffected in *Ring1* mutant MNs (**Fig. 7B,C**, **S4B**). These results indicate that assembly and maintenance of PHC2- and CBX2-containing aggregates in MNs are resistant to loss of RING1 proteins.

By contrast, the number and size of BMI1-containing puncta was markedly reduced in MNs of *Ring1* mutants (**Fig. 7B,C**). We considered whether the reduced detection of BMI1 protein reflected decreased *Bmi1* expression or compensatory changes in expression of its paralog *Pcgf2*. RNAseq analysis of MNs purified from *Ring1* mutants showed that *Bmi1* expression was not reduced, but instead was upregulated in *Ring1 MNΔ* mice, whereas *Pcgf2* expression was unchanged (**Fig. 7D**). We confirmed the presence of *Bmi1* mRNA in MNs by HCR **(Fig. S6A)**. However, overall BMI protein levels, as measured by immunofluorescence, were reduced in MNs of *Ring MNΔ* mice **(Fig. S6B)**, suggesting that RING1 loss affects BMI1 at the protein rather than transcriptional level.

These observations indicate that *Ring1* mutants, there is a loss of formation of BMI1- containing puncta, while CBX and PHC proteins retain the capacity to assemble into subnuclear clusters. Thus, the distinct *Hox* expression phenotypes in PRC2 and PRC1 mutants are associated with different degrees of PcG body loss, with BMI1-containing aggregates resistant to H3K27me3 depletion.

## DISCUSSION

Axial patterning depends on spatially restricted *Hox* gene expression, which underlies the specification of cell types along the rostrocaudal axis of bilaterians (Mann and Glassford, 2024, Smith and Kratsios, 2024, Wellik, 2024). While the roles of *Hox* genes in motor neuron columnar and pool fate specification are well established, less is known about the mechanisms that maintain *Hox* expression boundaries as neural progenitors exit the cell cycle and differentiate. Our findings demonstrate that PHC2-dependent *Hox* repression is critical during rostrocaudal patterning and in early postmitotic motor neurons. We found that disruption of PHC-dependent polymerization leads not only to caudal shifts in *Hox* expression but also the erosion of normally mutually exclusive *Hox* domains, resulting in defects in motor neuron specification. These observations highlight stable *Hox* repression as essential for preserving positional identities during spinal cord development, even after the dissipation of initial patterning cues.

### Early patterning of the rostrocaudal neuraxis

*Hox* genes are activated in overlapping collinear domains during early axial patterning of the vertebrate spinal cord. Expression is initiated by graded morphogens including retinoic acid (RA) and fibroblast growth factor (FGF), which preferentially induce rostral and caudal *Hox* paralogs, respectively (Miller and Dasen., 2024). The initial induction of *Hox* genes, and therefore early axial patterning of the embryo, appears to precede neurogenesis (Metzis et al., 2018). Opposing morphogen gradients establish presumptive *Hox* domains in neural progenitors, providing an initial framework for rostrocaudal identity. As development proceeds, early expression patterns are sharpened and stabilized through extensive cross-repressive interactions among *Hox* paralogs, ensuring the mutual exclusivity of Hox programs required for the development of segmentally-restricted MN fates (Philippidou and Dasen, 2013).

At the chromatin level, multiple layers of epigenetic regulation contribute to the establishment of *Hox* expression boundaries during early spinal cord patterning. Prior to neural differentiation, Polycomb repressive complex 2 (PRC2) deposits H3K27me3 across inactive *Hox* loci, creating a repressive chromatin landscape that restricts premature or ectopic transcription (Loubiere et al., 2019, Healy et al., 2019, Coleman and Struhl, 2017). Spatially controlled relief of this repression, driven by RA, FGF, Wnt, and TGFβ signaling, allows selective activation of appropriate *Hox* genes (Mazzoni et al., 2013, Mouilleau et al., 2021, Liu et al., 2001, Nordstrom et al., 2006). In parallel, chromatin architectural features—including topologically associating domains and boundary elements bound by CTCF—help insulate *Hox* clusters, limiting the spread of activating signals and reinforcing domain integrity (Narendra et al., 2016, Narendra et al., 2015). Together, these mechanisms ensure that early patterning cues are translated into stable, segment-specific transcriptional programs in neural progenitors.

Following MN differentiation, however, maintenance of *Hox* expression boundaries becomes uncoupled from the morphogen gradients that initially establish rostrocaudal identity. As progenitors exit the cell cycle and extrinsic patterning cues subside, *Hox* genes must remain durably repressed in appropriate segments to preserve motor neuron subtype identity. How this long-term repression is achieved in postmitotic neurons remains incompletely understood. Our results show that PHC2-dependent nuclear condensation is important for the restriction of *Hox* TFs during spinal MN differentiation.

Together, these observations indicate that stabilization of motor neuron subtype identity involves a developmental shift from signal-dependent and chromatin regulatory mechanisms in progenitors to a more persistent, structure-based modes of gene repression. This late regulatory phase may be especially important for motor circuit maturation, given emerging evidence that *Hox* genes continue to function in early postnatal stages (Feng et al., 2022, Catela et al., 2022, Kc et al., 2024).

### PHC2 assemblies are required for stable repression of Hox TFs in spinal MNs

Our findings indicate that PRC1-mediated repression of *Hox* genes is reinforced, rather than relaxed, as neural progenitors differentiate into postmitotic motor neurons. In neural progenitors, BMI1, PHC2, and CBX2 exhibit relatively diffuse nuclear distributions, whereas in postmitotic brachial motor neurons these same proteins assemble into larger, discrete subnuclear assemblies. The progressive increase in the size of PRC1 assemblies during differentiation suggests that Polycomb repression is structurally stabilized as cells exit the cell cycle, consistent with the increased requirement to maintain transcriptional memory in postmitotic neurons.

Polymerization of Polyhomeotic proteins is thought to stabilize and propagate Polycomb repressive domain, thus contributing to chromatin compaction (Isono et al., 2013, Kim et al., 2002). We found disruption of PHC2-dependent polymerization uncouples PRC1 expression from its repressive function, revealing a role for PRC1 assembly architecture in transcriptional silencing. Expression of dominant-negative PHC2 resulted in a reduction in PRC1 assembly size without affecting overall PRC1 protein abundance or generic motor neuron identity. Notably, the extent of *Hox* derepression closely tracked the degree of polymer disruption within individual nuclei, with smaller and more diffuse BMI1 foci associated with higher levels of ectopic HOXC9 expression. These results argue that PRC1 assemblies are not passive accumulations of Polycomb proteins but instead function as structural units whose integrity is required for effective repression of *Hox* loci.

Comparison of PRC2 and PRC1 loss-of-function mutants further clarifies the relationship between Polycomb architecture and *Hox* repression in motor neurons. Motor neuron–restricted deletion of *Ezh1/2* resulted in efficient depletion of H3K27me3 but did not disrupt PRC1 subnuclear organization, as PHC2- and CBX2-containing assemblies remained intact and BMI1 puncta were modestly increased. In contrast, loss of the core PRC1 component *Ring1* caused a selective loss of BMI1-containing assemblies, coinciding with *Hox* dysregulation and loss of Hox-dependent motor neuron identities. Together with the *dnPhc2* experiments, these findings indicate that PRC1 assembly architecture constitutes a critical repressive mechanism maintaining *Hox* profiles throughout motor neuron maturation.

Importantly, PHC-dependent PRC1 assemblies remain necessary after motor neuron differentiation. Targeting *dnPhc2* expression specifically to postmitotic motor neurons under the *Hb9* promoter was sufficient to reduce the size of PRC1 assemblies and induce *Hox* derepression. This finding indicates that PRC1-mediated repression is continuously required throughout neural differentiation and maturation, rather than being locked in during progenitor stages. Together, these observations support a model in which PRC1 assemblies provide a stable repressive architecture that safeguards *Hox* profiles as neural cells transition from proliferative progenitors to terminally differentiated motor neurons.

### PRC1 assemblies are required for Hox-dependent motor neuron differentiation

Disruption of PRC1 assembly integrity has direct consequences for the differentiation of spinal motor neuron subtypes. Following expression of *dnPhc2*, misexpression of caudal *Hox* genes causes a failure to specify Hox-dependent motor columns, including the lateral motor column (LMC) and preganglionic motor column (PGC). This was reflected by a pronounced loss of FOXP1 expression, together with reduced expression of columnar-specific markers such as RALDH2 and pSMAD. These defects occurred without affecting generic motor neuron markers, indicating that PRC1 assembly disruption selectively compromises subtype differentiation rather than motor neuron identity itself. The resulting phenotypes closely resemble those observed in *Ring1* mutants, underscoring the requirement for intact PRC1 architecture in establishing Hox-dependent neuronal identities.

In contrast, motor neuron subtypes specified independently of *Hox* activity were preserved or expanded following disruption of PHC2-dependent PRC1 assembly. In *dnPhc2*-expressing embryos, motor neurons that failed to adopt LMC or PGC identities instead exhibited molecular features of the medial motor column, as indicated by increased expression of MECOM and LHX3. This shift toward MMC identity differs from *Ring1* mutants, where motor neurons predominantly acquire an HMC identity (Sawai et al., 2022). While the mechanisms accounting for this difference are unclear, they may reflect distinct roles of RING and PHC proteins, or varying degrees of residual repressive activity. Collectively, these fate transformations indicate that sustained Polycomb-mediated repression is required to prevent motor neurons from defaulting toward ancestral *Hox*-independent identities.

An important next step will be to determine how PRC1 assemblies engage chromatin to enforce long-term *Hox* repression in differentiated neurons, including examining whether assembly formation directly constrains three-dimensional genome organization or operates through selective stabilization of repressive chromatin contacts. It will also be informative to determine whether similar Polycomb-dependent mechanisms maintain positional identity in other postmitotic lineages, and explore whether dynamic changes in PRC1 subunit compositions contribute to fate specification. Finally, dissecting how distinct PRC1 subassemblies contribute to repression may reveal how Polycomb complexes achieve both stability and flexibility during development.

## MATERIALS AND METHODS

### Animal lines and embryo collection

All animal protocols were approved by the Institutional Animal Care and Use Committee (IACUC) at NYU Langone Health. Timed matings for mouse embryos were performed by observing a vaginal plug and designated as embryonic day (E) 0.5. Embryos were harvested at E12.5 and fixed in 4% paraformaldehyde for 90 minutes at 4°C.

*Ezh1 fl*ox/flox (Hidalgo et al., 2012), *Ezh2 flox/flox* (Su et al., 2003), and *Olig2-Cre* (Dessaud et al., 2007) mouse lines have been previously described. Generation of *EzhMNΔ* mice was performed by crossing *Ezh1flox/flox, Ezh2flox/flox* and *Olig2-Cre* mice. Generation of *Ring1MNΔ* mice was performed by crossing *Ring1-/-::Rnf2flox/flox* (Cales et al., 2008, del Mar Lorente et al., 2000) and *Olig2Cre* mice.

### DNA constructs

The *pCIG* expression vector was used for electroporation-based expression in chick embryos and contains a CAG promoter and an internal ribosome entry site (IRES) driving nuclear GFP expression. The following constructs were used: *pCIG* (control), *pCIG-Phc2*, and *pCIG-dnPhc2*. The dominant-negative *Phc2* (*dnPhc2*) construct contains a single L307R missense mutation within the End-Loop-Helix (ELH) domain, which disrupts Sterile Alpha Motif (SAM) domain–mediated polymerization of PHC2 and thereby inhibits canonical PRC1 assembly formation.

For post-mitotic motor neuron–restricted expression, constructs were cloned downstream of the *Hb9* promoter. These included *Hb9-GFP*, *Hb9-PHC2-GFP*, and *Hb9- dnPhc2-GFP*. GFP expression was used to identify transgene-expressing motor neurons in all experiments. All plasmids were sequence verified prior to use.

### *In ovo* electroporation

*In ovo* electroporation of chick embryos was performed at Hamburger & Hamilton (HH) stage 13-15 and analyzed at HH25-31. Fertilized chicken eggs (sourced from Charles River) were incubated at 39.5°C in a humidified chamber for 48 hours until HH13-14.

The top of the eggshell was removed and 1-3 µg/µl DNA (500-2500 ng/µL expression plasmid and pBKS carrier DNA) containing ∼0.02% Fast Green was injected into the central canal of the neural tube using a sharpened glass capillary tube. Electrodes (Platinum/Iridium (80%/20%), 250 μm diameter, UEPMGBVNXNND, FHC Inc) were placed on both sides of the neural tube (4 mm separation) and DNA was electroporated using an ECM 830 electroporator (ECM 830, BTX; 25 V, 4 pulses, 50ms duration, 1 s interval). Eggs were sealed with parafilm and incubated for 48-72 hr prior to fixation.

Unilateral electroporation enabled direct comparison between electroporated (GFP- positive) and contralateral non-electroporated (GFP-negative) cells within the same embryo. Results shown in figures are representative of at least three electroporated embryos from two or more experiments in which electroporation efficiency in MNs was above 60%.

### Slide immunohistochemistry

Embryos were fixed in 4% PFA for 1.5 hr at 4°C, washed 5–6 times in cold PBS for 15– 30 minutes each wash, and cyroprotected overnight in 30% sucrose. Tissue was embedded in OCT, frozen in dry ice, and sectioned at 16 µm on a cryostat. For antibody staining, slides were placed in PBS for 5 min to remove OCT. Sections were then transferred to humidified trays and blocked for 20–30 min in 0.75 mL/slide of PBT (PBS, 0.1% Triton) containing 1% Bovine serum albumin (BSA). The blocking solution was replaced with primary staining solution containing antibodies diluted in PBT with 0.1% BSA. Primary antibody staining was performed overnight at 4°C. Slides were then washed three times for 5 min each in PBT. Fluorophore-conjugated secondary antibodies were diluted 1:1000 in PBT and filtered through a 0.2 µm syringe filter.

Secondary antibody solution was added to slides (0.75 ml/slide) and incubated at room temperature for 1 hr. Slides were washed three times in PBT, followed by a final wash in PBS. Coverslips were placed on slides using 110 µl of Vectashield (Vector Laboratories).

Antibodies against Hox proteins and MN subtype markers were generated as described (Dasen et al., 2008, Dasen et al., 2003, Dasen et al., 2005). Additional antibodies were used as follows: rabbit anti-RNF2 (RING1B) (Abcam, 1:5000), rabbit anti-CBX2 (Bethyl, 1:5000), rabbit anti-H3K27me3 (Cell Signaling, 1:2000), rabbit anti-BMI1 (Bethyl, 1:1000), mouse anti-BMI1 (NSJ Bioreagents, 1:1000) and rabbit anti-PHC2 (EMD Millipore, 1:2000). Detailed description of antibodies is provided in Table S1.

### Hybridization chain reaction (HCR) RNA-FISH

HCR was performed on fresh-frozen tissue sections following the third-generation protocol (v3.0) from Molecular Instruments (Choi et al., 2018). DNA probe sets, HCR amplifiers, and associated buffers were obtained from Molecular Technologies (Beckman Institute, Caltech). Custom probes were synthesized by Integrated DNA Technologies using oPools Oligo Pools. Slides were mounted using 110 µl Vectashield (Vector Laboratories), and confocal images were acquired on a Zeiss LSM700 microscope. Probes with indicated amplifier sequences were as follows: B1: mouse *Bmi1*, chick *Hoxc9*; B2: mouse *Hb9* (*Mnx2*), chick *Hoxc10*. Probe sequences are available from the Lead author upon request.

### Confocal and Airyscan imaging

Fluorescent imaging was performed using laser-scanning confocal microscopy and Airyscan super-resolution microscopy using the NYU Langone’s Microscopy Laboratory (RRID: SCR_017934). Confocal imaging was used for low-magnification survey images and qualitative assessment of marker expression, while Airyscan imaging was used for high-resolution visualization and quantitative analysis of PRC1 assembly organization, as previously described (Huff., 2015).

All measurements were performed on a Zeiss LSM 880 microscope equipped with the AiryScan detector, an Argon laser (Melles-Griot) for 488 nm excitation and a Zeiss Plan-Apochromat 63×/1.4 NA DIC M27 Oil objective. Raw Airyscan data were processed using standard Airyscan reconstruction algorithms to improve spatial resolution and signal-to-noise ratio relative to conventional confocal imaging. Identical acquisition settings and reconstruction parameters were applied across experimental conditions within each dataset.

For all quantitative analyses, imaging was performed under non-saturating conditions, and detector gain, laser power, and pinhole settings were held constant between control and experimental samples. Representative images shown in figures were selected from datasets used for quantification.

### PRC1 assembly segmentation and size quantification

PRC1 assemblies were identified and quantified from Airyscan super-resolution images based on immunofluorescent signal from PRC1 components, including BMI1, PHC2, RING1B, or CBX2, as indicated for each experiment. Quantitative analyses were performed on single optical sections or maximum-intensity projections encompassing individual motor neuron nuclei.

Images were preprocessed uniformly across datasets to reduce background signal prior to segmentation. Nuclear regions of interest were defined based on DAPI staining and motor neuron markers, and only nuclei with clearly resolved PRC1 signal were included in subsequent analyses. PRC1 assemblies were segmented using intensity-based thresholding followed by particle analysis to identify discrete polymer structures.

For each segmented polymer, area measurements were extracted and converted to square micrometers (µm²). Polymers were categorized by size into three bins: small (<0.04 µm²), medium (0.04–0.08 µm²), and large (>0.08 µm²). Polymer size distributions were quantified on a per-nucleus basis and pooled across embryos for each condition.

All segmentation parameters and size thresholds were applied identically across control and experimental groups within each experiment. Analyses were performed on images acquired using identical imaging and reconstruction settings. Data was collected from multiple independent embryos, as indicated in the figure legends.

### Quantification and statistical analysis

Image-based quantifications were performed using standardized analysis pipelines applied uniformly across experimental conditions. PRC1 assembly size measurements, polymer size distributions, and colocalization analyses were conducted on a per-nucleus basis and pooled across multiple independent embryos. For all statistical analyses, three biological replicates were used, with each replicate corresponding to an independent embryo.

Statistical analyses were performed using appropriate tests as indicated in the figure legends. Data are presented as mean values with measures of variability as specified. No statistical methods were used to predetermine sample size. All analyses were performed using standard image analysis and statistical software.

### Artificial Intelligence tools

ChatGTP was used to correct grammar and typos.

## Acknowledgements

We thank Timothee Lionnet, Estaban Mazzoni, and Jessica Treisman for their critical feedback on the work. We thank NYU Langone’s Microscopy Laboratory facility for assistance with high resolution imaging. Parts of the text and figures for the manuscript are reproduced from the PhD thesis of Alexander Miller (New York University, 2026).

## Author contributions

Conceptualization: A.K.M., J.S.D.; Data curation: A.K.M.; Formal analysis: A.K.M., J.S.D.; Funding acquisition: A.K.M., J.S.D.; Investigation: A.K.M., A. S.-F., A.E.C.; Methodology: A.K.M., A. S.-F., A.E.C.; Project administration: A.K.M.; Resources: J.S.D.; Supervision: J.S.D.; Validation: A.K.M., J.S.D.; Visualization: A.K.M.; Writing – original draft: A.K.M, J.S.D.; Writing – review & editing: A.K.M, J.S.D..

## Data availability

Original data is available from the authors upon request.

