## Supplemental Figures S1-S6 for "PHC2-Dependent PRC1 Assemblies Stabilize Motor Neuron *Hox* Codes"

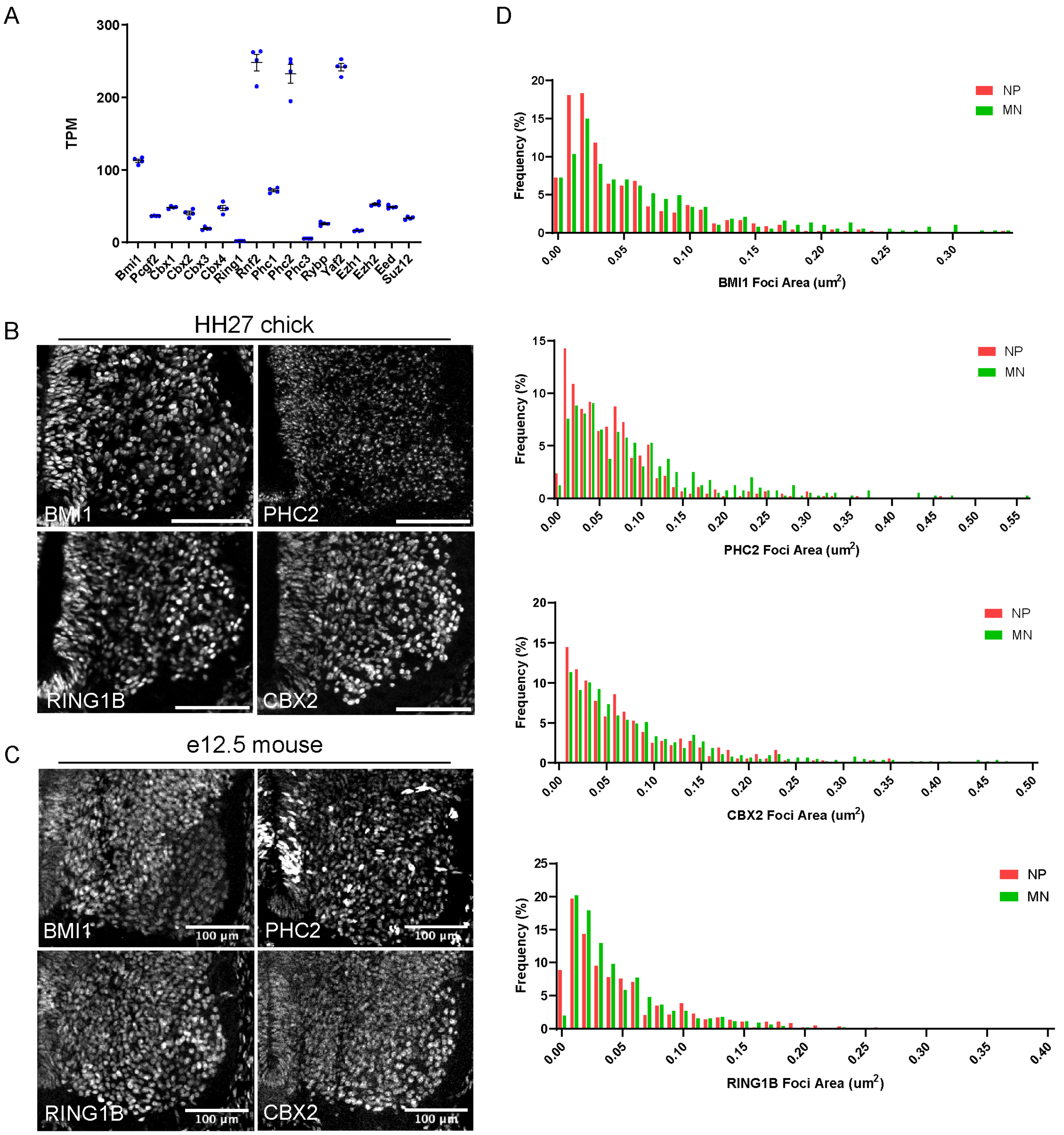


**Fig. S1. Expression of canonical PRC1 subunits during motor neuron differentiation.**
(A) Bulk RNA-seq data showing expression levels (transcripts per million [TPM]) of PRC1 and PRC2 subunits. Data points are from n=4 embryos, 3 sections/embryo.
(B) Expression of PRC1 protein subunits (BMI1, PHC2, RING1B, CBX2) in brachial HH27 chick spinal cord. Scale bars, 100 µm.
(C) Expression of PRC1 components in brachial spinal cord sections from E12.5 mouse embryos.
(D) Full distribution of PRC1 puncta sizes in NPs and MNs corresponding to Airyscan datasets in Fig. 1A-D.

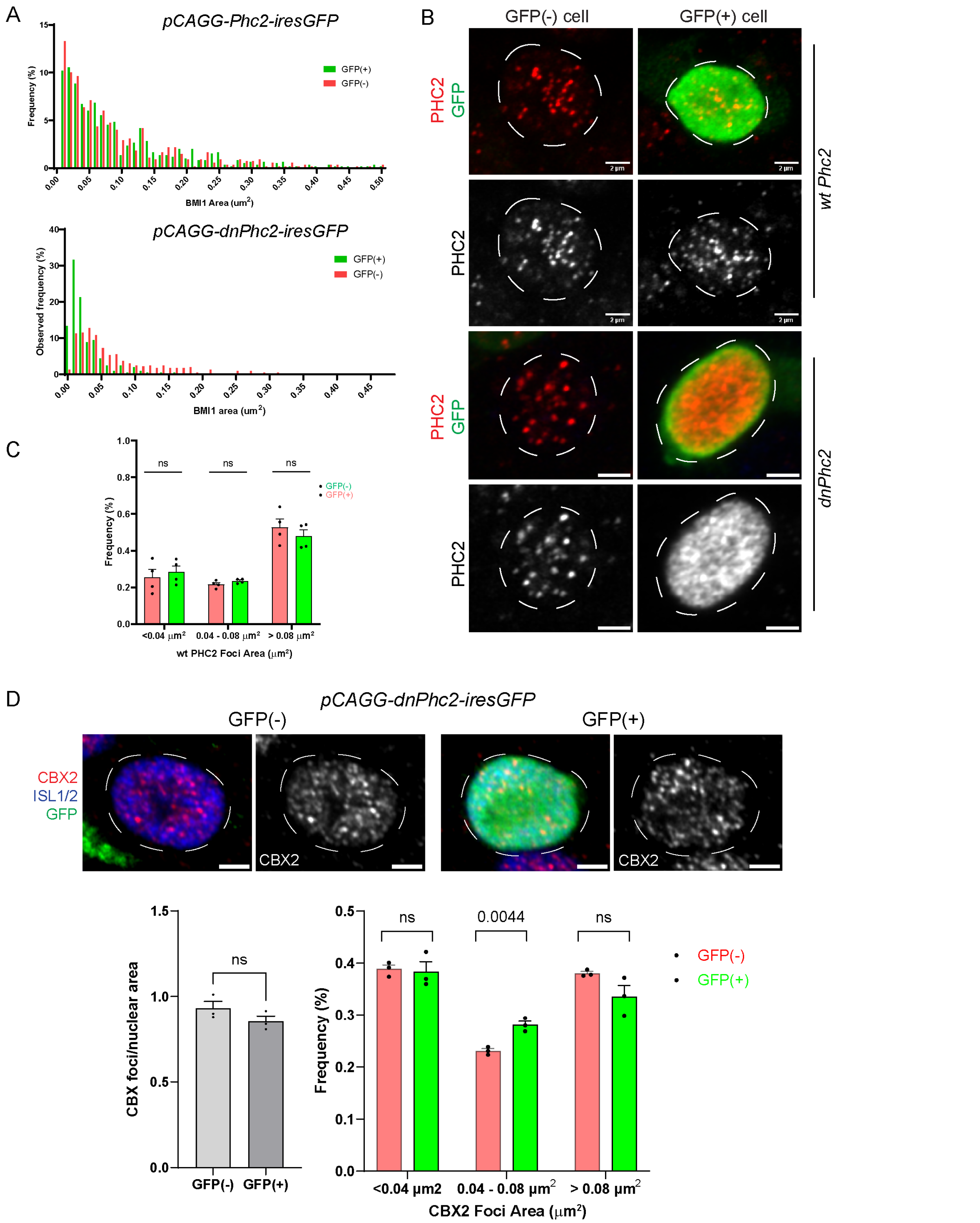


**Fig. S2**: **Effects of blocking PHC polymerization in MNs.**

(A) Full size distributions of BMI1-containing puncta after *Phc2* and *dnPhc2* expression.

(B) Analyses of PHC2 localization after *Phc2* and *dnPhc2* expression. PHC2 becomes more diffuse in *dnPhc2* expressing neurons. Scale bars, 2 µm.

(C) Distribution of PHC2 puncta sizes in GFP(-) and GFP(+) nuclei in wt *Phc2*-expressing neurons.

(D) Expression of *dnPhc2* does not disrupt formation of CBX2-containing puncta. Quantification of the number of CBX foci/area and frequencies of large and small foci are shown. Bar graphs show mean ± SEM for n=3 GFP (-) control and GFP (+) *dnPhc2* embryos, 3 sections/embryo. Scale bars, 2 µm.

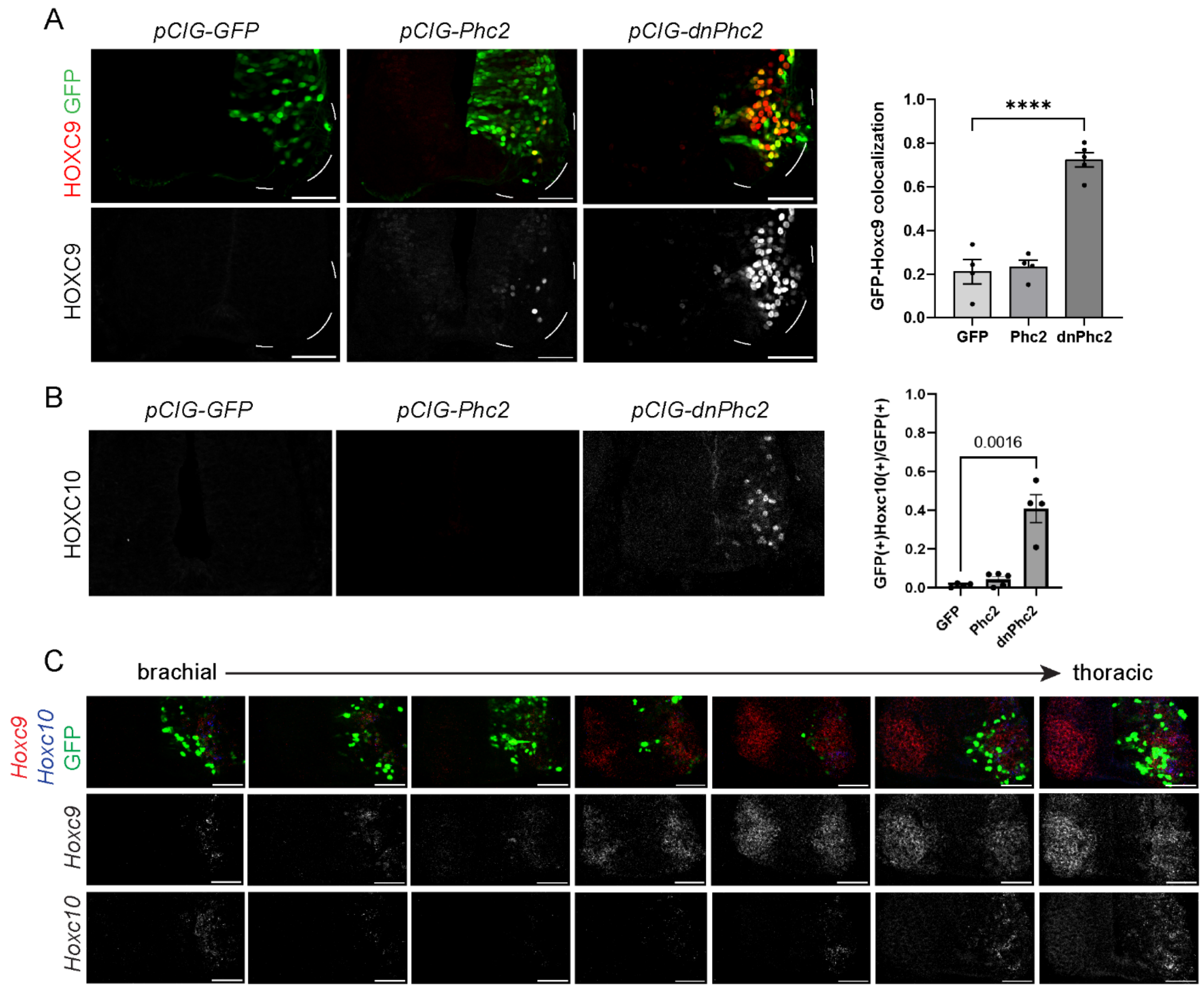


**Fig. S3. Expression of Hox transcription factors after expression of *GFP*, *Phc2* and *dnPhc2*.**
(A) Expression of HOXC9 in brachial spinal cord of embryos electroporated with *pCIG-GFP* (control), *pCIG-Phc2*, or *pCIG-dnPhc2*. Bar graphs show mean fractions ± SEM for n=4-5 embryos, 3 sections/embryo. ****$p < 0.0001$. Scale bars, 50 µm
(B) Expression of HOXC10 in brachial spinal cord of embryos electroporated with *pCIG-GFP* (control), *pCIG-Phc2*, or *pCIG-dnPhc2*. Bar graphs show mean fractions ± SEM for n=4-5 embryos, 3 sections/embryo.
(C) HCR *in situ* hybridization showing ectopic expression of *Hoxc9* and *Hoxd10* in *pCIG-dnPhc2* electroporated embryos. Scale bars, 50 µm.

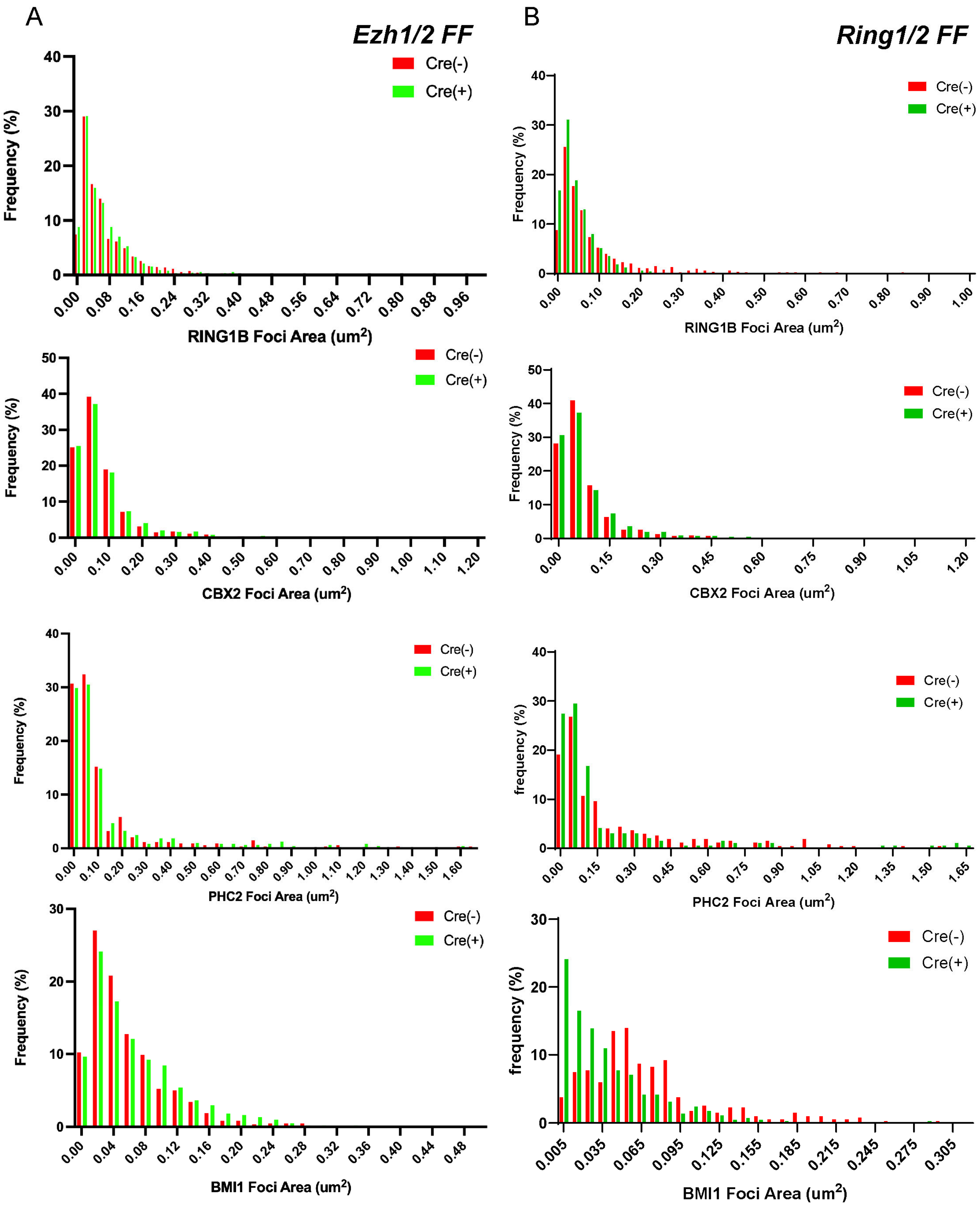


**Fig. S4. Full distributions of PRC1 assembly sizes in *Ezh1/2* and *Ring1* mutant MNs.**
(A) Distributions of RING1B, CBX2, PHC2, and BMI1-containing puncta in control (Cre-) and *Ezh* mutant (Cre+) MNs.
(B) Distributions of RING1B, CBX2, PHC2, and BMI1-containing puncta in control (Cre-) and *Ring1* mutant (Cre+) MNs.

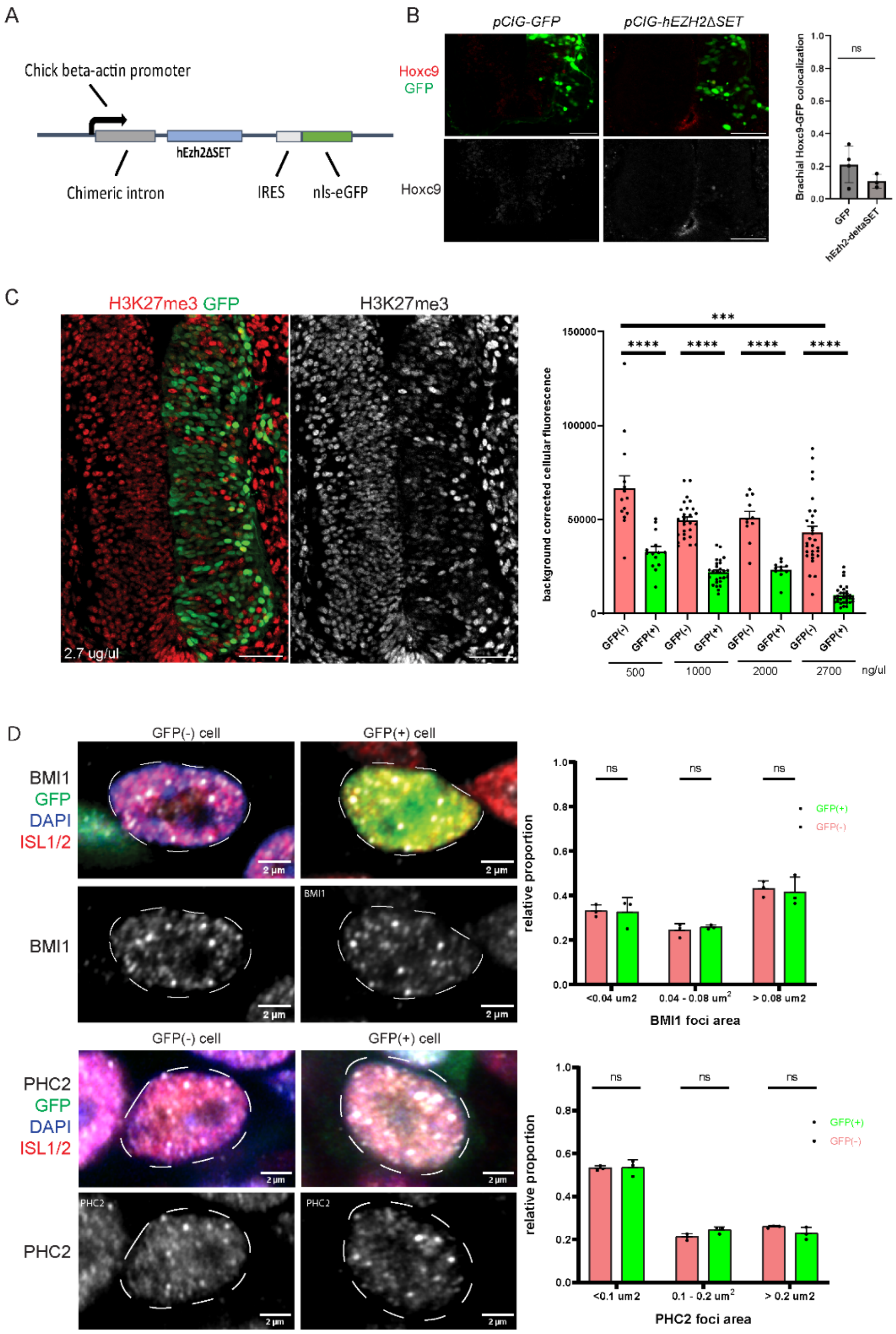


**Fig. S5. PRC1 assemblies are resistant to reduction in H3K27me3.**
(A) Schematic of the *pCIG-hEzh2ΔSET* construct.
(B) HOXC9 is not derepressed after *pCIG-hEzh2ΔSET* expression. Right side of the spinal cord is electroporated. Quantification of GFP:HOXC9 colocalization shown on right. Bar graphs show mean fractions ± SEM for n=3 embryos, 3 sections/embryo. Scale bars, 50 µm.
(C) Representative images of H3K27me3 following expression of *hEzh2-ΔSET*. Corrected signal intensity across increasing injection concentrations shown on right. Bar graphs show mean ± SD. Individual sections are plotted. Scale bars, 50 µm.
(D). High-resolution imaging of BMI1 and PHC2 puncta in embryos electroporated with 2.7µg/µl *pCIG-hEzh2ΔSET*. Right panels show the distribution of BMI1 and PHC2 polymer sizes binned as small (<0.04 µm$^2$), medium (0.04-0.08 µm$^2$), and large (>0.08 µm$^2$). Bar graphs show mean ± SD, from n=3 embryos, 3 sections/embryo.

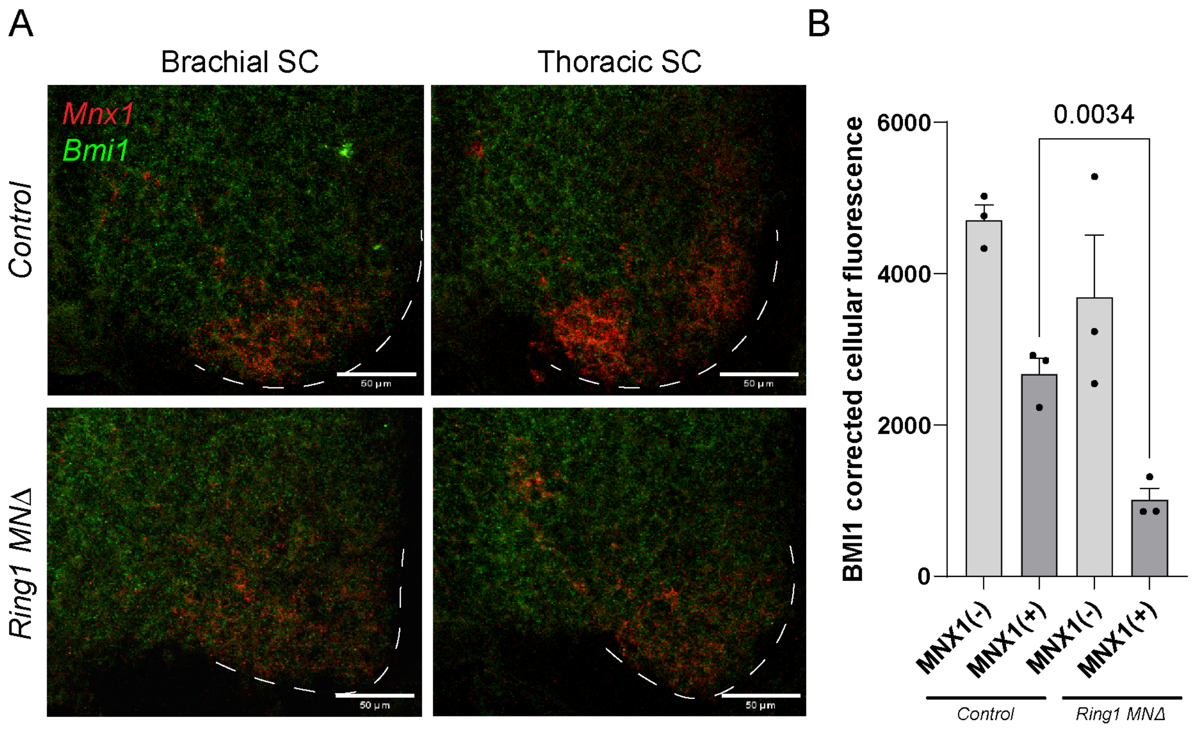


**Fig. S6. Loss of *Ring1* genes leads to a reduction in BMI protein expression.**
(A) HCR in situ hybridization showing expression of *Bmi1* in motor neurons (marked by *Mnx1* expression) in control and *Ring1* mutants in brachial and thoracic segments.
(B) Quantification of BMI1 immunofluorescence of MNX1(+) (also known as HB9) MNs in control and *Ring1* mutants. Three embryos were analyzed. Bar graphs show mean ± SEM for n=3 embryos, 3 brachial sections/embryo.

**Table S1. Antibodies used in this study.**
